# Circuit-Specific Targeting of Astrocytes for Genetic Control

**DOI:** 10.1101/2022.03.07.483355

**Authors:** Alyssa Thompson, Rachel Arano, Katarina Ramos, Yerim Kim, Ying Li, Wei Xu

**Affiliations:** Department of Neuroscience, University of Texas Southwestern Medical Center, Dallas, TX 75390, USA

## Abstract

Astrocytes are integral functional components of brain circuits. They ensheath the connections between neurons to form tripartite synapses. They react to local neuronal activities and release signaling molecules to regulate synaptic transmission. Their dysfunctions impair synaptic functions and are implicated in neuropsychiatric disorders. Increasing evidence indicates that astrocytes are diverse and they have distinct features and functions in different circuits. However, selectively targeting and controlling astrocytes in a circuit-specific manner is technically challenging. Recently we constructed a series of anterograde transneuronal viral vectors based on the yellow fever vaccine YFV-17D. These YFV-17D derivatives express fluorescent proteins almost exclusively in neurons. However, we find that YFV-17D carrying DNA recombinase Cre infect astrocytes associated with the traced neuronal pathways and express Cre to turn on reporter genes. The targeting of astrocytes is at a whole-brain level but specific to the neuronal circuits traced. Therefore, YFV-17D vectors carrying DNA recombinases provide tools for selectively and genetically targeting pathway-specific astrocytes. This new technology will also allow us to reveal the roles of astrocytes in specific neuronal circuits in normal brain functions and diseases.

## Introduction

Astrocytes are one of the most abundant cell types in the brain. They provide structural, nutritional and metabolic support to neurons. Equally important, they are an essential component of synaptic communication in the brain. Structurally, astrocytic processes are one of three elements for tripartite synapses. Electron microscopic reconstructions of the synapses in the brain reveal that over 50-95% of the synapses are contacted or ensheathed by astrocytic processes (Chai et al., 2017; Xu-Friedman et al., 2001). Astrocytes also contribute to synapse formation and maintenance. Synapses are formed primarily after astrogenesis during development, and the molecules expressed or released from astrocytes determine many synaptic features (Farhy-Tselnicker and Allen, 2018). A few traditional neuronal adhesion molecules are expressed by astrocytes for synapse maintenance (Stogsdill et al., 2017). Finally, bidirectional communication exists between neurons and astrocytes in synaptic transmission. Astrocytes are excitable cells and they react to neurotransmitters released by synaptic terminals with calcium transients generated locally in their processes, which can spread to the soma or even neighboring astrocytes (Perea and Araque, 2005; Perea et al., 2009). The calcium activity may trigger the release of molecules such as glutamate, D-serine, adenosine and ATP, which in turn modulates synaptic transmission in the neuronal part of the synapse (Savtchouk and Volterra, 2018). This bidirectional interaction may regulate the strength and plasticity of synaptic transmission or even neuronal firing (Deemyad et al., 2018).

Astrocytes tile the whole brain. Despite the “even” distribution, astrocytes are highly diverse (Khakh and Sofroniew, 2015). A systematic comparison of the astrocytes in the hippocampus and the striatum reveals distinct features of these cells in morphology, molecular profiles, and reactions to signal molecules (Chai et al., 2017). Single-cell analysis of cell lineage, genetic identity and reaction to inflammatory stimulus reveals seven to ten subtypes of astrocytes (Bandler et al., 2022; Hasel et al., 2021). On top of the diversity of the astrocytes themselves, astrocytes may exert distinct influences on synapses in the different circuits even if the synapses are located in the same brain region. For example, in the striatum, the subpopulations of astrocytes differentially regulate the synaptic transmission of direct vs. indirect striatal pathways (Martin et al., 2015). These studies all indicate that the functions of the astrocytes are closely coupled to the synapses of the specific neuronal circuits they are associated with.

Revealing the circuit-specific roles of astrocytes relies on technologies which target and control astrocytes in a circuit-specific manner. Recent research on astrocytes has been facilitated by the availability of many genetic or viral tools to selectively target astrocytes. Combining these tools allows us to monitor or manipulate brain region-specific astrocytes. However, it is still not possible to separate the astrocytes distributed in the same region that are associated with distinct neuronal pathways. In our efforts to develop novel transneuronal viral vectors based on yellow fever vaccine YFV-17D, we surprisingly noticed that the engineered YFV-17D carrying DNA recombinase Cre could effectively infect astrocytes associated with the traced neuronal circuits and turn on report genes. This therefore provides a technology to achieve circuit-specific genetic control of astrocytes at unprecedented precision.

## Results

We first examined the spreading of the replication-deficient anterograde transneuronal tracer virus, YFV^ΔNS1^-Cre (Li et al., 2021), in the brain of reporter mice--Ai9 mice expressing red fluorescent protein tdTomato in Cre-expressing cells. Deletion of NS1 makes YFV17D unable to replicate, therefore we must provide NS1 at the pre-synaptic sites for YFV to replicate and propagate. We injected adeno-associated viruses (AAVs) mediating the inducible expression of YFV-17D gene NS1, including AAV-tTA and AAV-TRE-NS1, into the motor cortex of Ai9 mice. The mice were fed with a diet containing doxycycline (Dox) as described previously to restrict the replication and spreading of YFV^ΔNS1^-Cre to a short time window. Two weeks later we injected YFV^ΔNS1^-Cre into the same locus. After another 8-10 days, we fixed the brains for histological analysis. In our previous work with YFV^ΔNS1^-Cre in combination with AAV reporter (eg. AAV-DIO-jGCaMP7f), the NS1 gene needed to be expressed in the post-synaptic neurons to allow of YFV^ΔNS1^ to reach the postsynaptic neurons to efficiently turn on reporter genes like jGCaMP7f (when AAV-DIO-jGCaMP7f was the reporter system). However, to our surprise, in Ai9 mice, without providing the NS1 gene in the downstream regions, we observed a large number of cells expressing tdTomato throughout the brain (Fig. 1a). These tdTomato-positive cells were in brain regions that were known downstream targets of the neurons in the motor cortex, such as the contralateral cortex, the ipsilateral and contra lateral dorsal striatum, the dorsal thalamus, zona incerta, pontine nucleus and brain stem structures. These traced cells were preferentially distributed in the hemisphere ipsilateral to the injection site.

**Fig.1.**
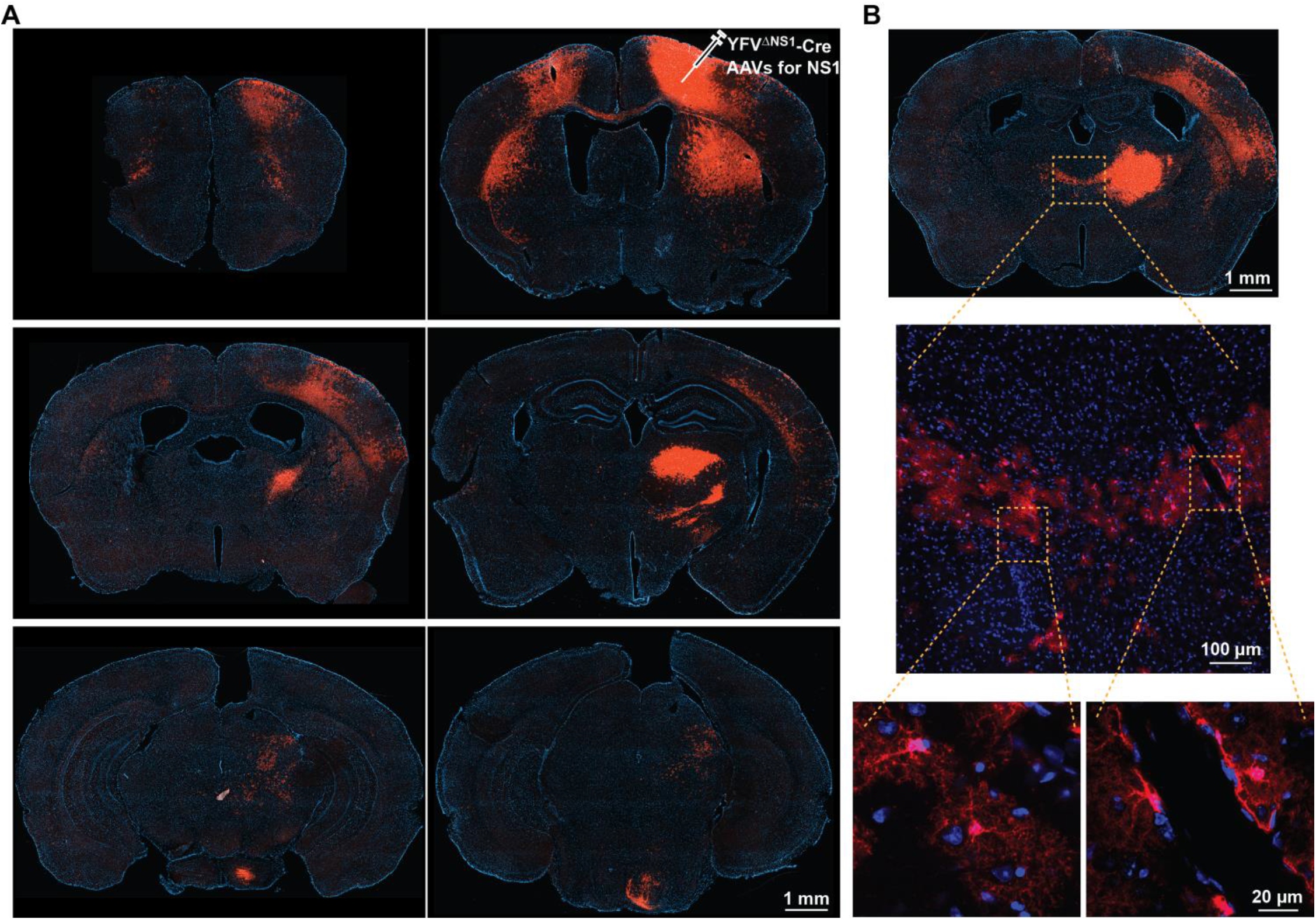
Locally injected YFV^ΔNS1^-Cre turns on reporter genes at the whole-brain level. (A) The expression of tdTomato (red) in the brain. The mouse had received an AAV injection (AAV-tTA and AAV-TRE-NS1) at the motor cortex (Mo) 22 days earlier and YFV^ΔNS1^-Cre injection in the same locus 8 days earlier. The blue color is the counterstaining with DAPI. (B) High-resolution photos of the tdTomato-positive cells.

A closer look demonstrated that most of the tdTomato-positive cells (except those at the viral injection site where NS1 was expressed) lacked the typical morphology of neurons (Fig. 1B). With dense processes and cloud-like morphology, they looked like astrocytes. Sometimes, these processes lined the walls of the brain blood vessels, consistent with the features of astrocytes.

To confirm the identities of these tdTomato-positive cells, we conducted immunostaining of markers for the major cell types in the brain (Fig.2). As expected, only a small fraction of these cells was positive for the neuronal marker NeuN. However, most of the cells were positive for a marker for astrocytes--S100β. These cells were not positive for a marker for oligodendrocyte precursor cells--NG2, or a marker for microglia--TMEM119, further supporting the identification of these cells as astrocytes.

**Fig.2.**
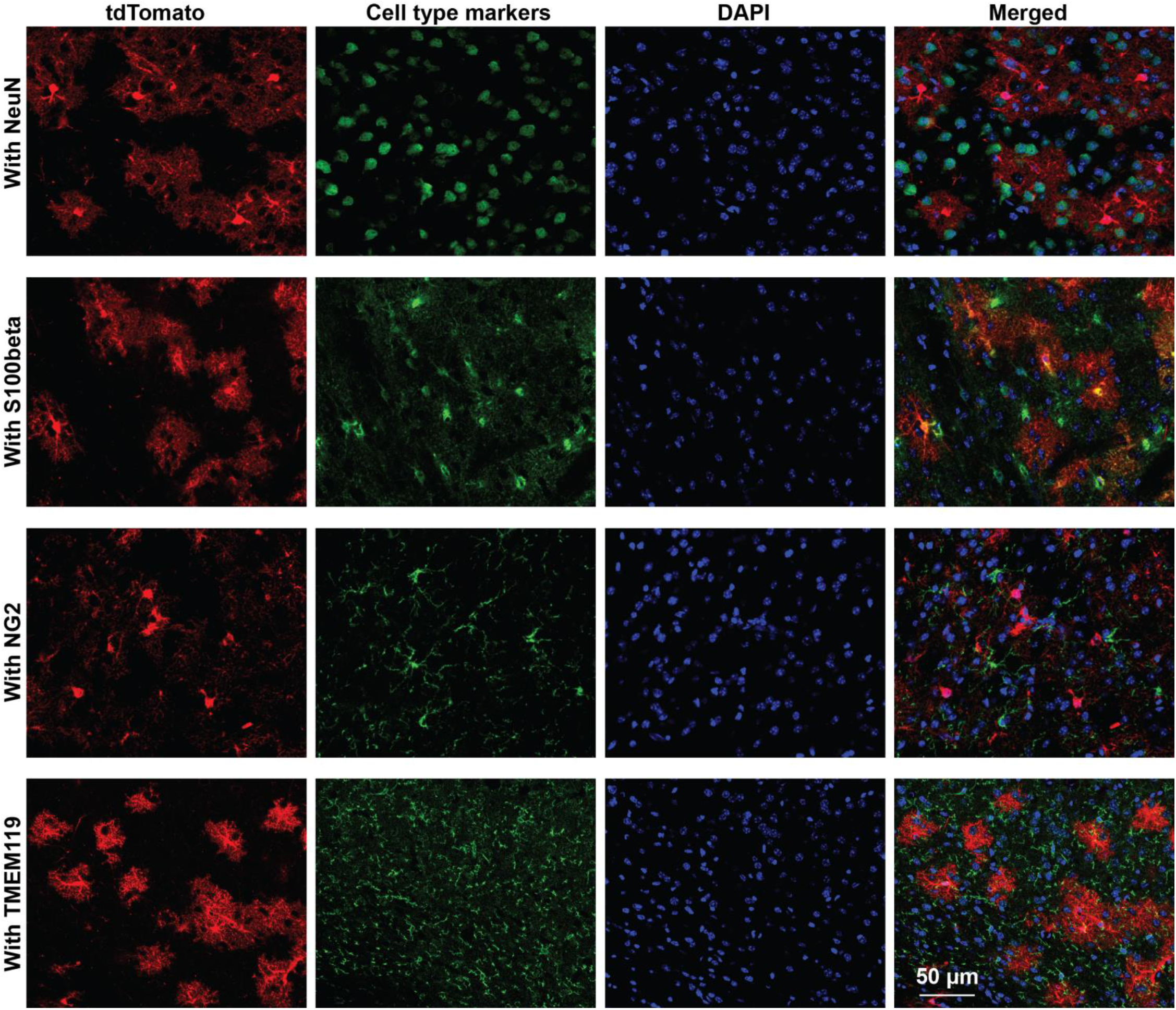
YFV^ΔNS1^-Cre turns on reporter genes in astrocytes. Brain sections from mice receiving YFV^ΔNS1^-Cre injection in the cortex immunostained with antibodies for the markers of different cells types, including NeuN for neurons, S100β for astrocytes, NG2 for oligodendrocyte precursor cells, and TMEM119 for microglia. The blue color is counterstaining with DAPI.

Next, we examined if the labeling was specific to Ai9 mice. We conducted the same viral injections in Ai6 mice that express green fluorescent protein ZSGeen1 in Cre-expressing cells. Similarly, we observed labeling of astrocytes in the regions receiving synaptic inputs form the viral injection site (Fig.2), suggesting that YFV^ΔNS1^-Cre could turn on Cre-dependent gene expression in different mouse lines. We also tested if YFV^ΔNS1^-Cre could turn on reporter genes carried by AAVs. We generated AAV5 expressing hM3Dq fused with mCherry under the control of an astrocyte-specific GfaABC1D promoter and injected this to the striatum (Chai et al., 2017). To our disappointment, we only detected a small number of mCherry-positive astrocytes and more mCherry was expressed in neurons (data not shown). The results indicated that YFV^ΔNS1^-Cre could not efficiently turn on an AAV reporter in the post-synaptic brain regions if NS1 was not provided. A possible reason is that the AAV infection triggered the defense mechanisms of astrocytes, which made these astrocytes resistant to YFV-17D infection.

Then we examined to what degree the astrocytes labeled by YFV^ΔNS1^-Cre were confined to the neuronal circuits YFV^ΔNS1^-Cre traveled along. We compared the distribution of the traced astrocytes in well-delineated circuits. We first took advantage of the projections from two neighboring cortical regions, the prefrontal cortex (PFC) and the motor cortex (Mo). These two cortical regions are known to selectively project to two distinct anatomical domains of the dorsal striatum--the dorsomedial striatum (DMS) and the dorsolateral (DLS), and through DMS/DLS regulate distinct behavioral modes, respectively (Fig. 3A). We injected into the PFC or Mo, respectively, AAVs expressing NS1 and then YFV^ΔNS1^-Cre. When YFV^ΔNS1^-Cre was injected into the PFC the traced astrocytes were primarily in the DMS (Fig. 3B); but when YFV^ΔNS1^-Cre was injected into the Mo, the traced astrocytes were in the DLS (Fig. 3C). This suggests that YFV^ΔNS1^-Cre only infected astrocytes associated with the neuronal circuits it traced.

**Fig.3.**
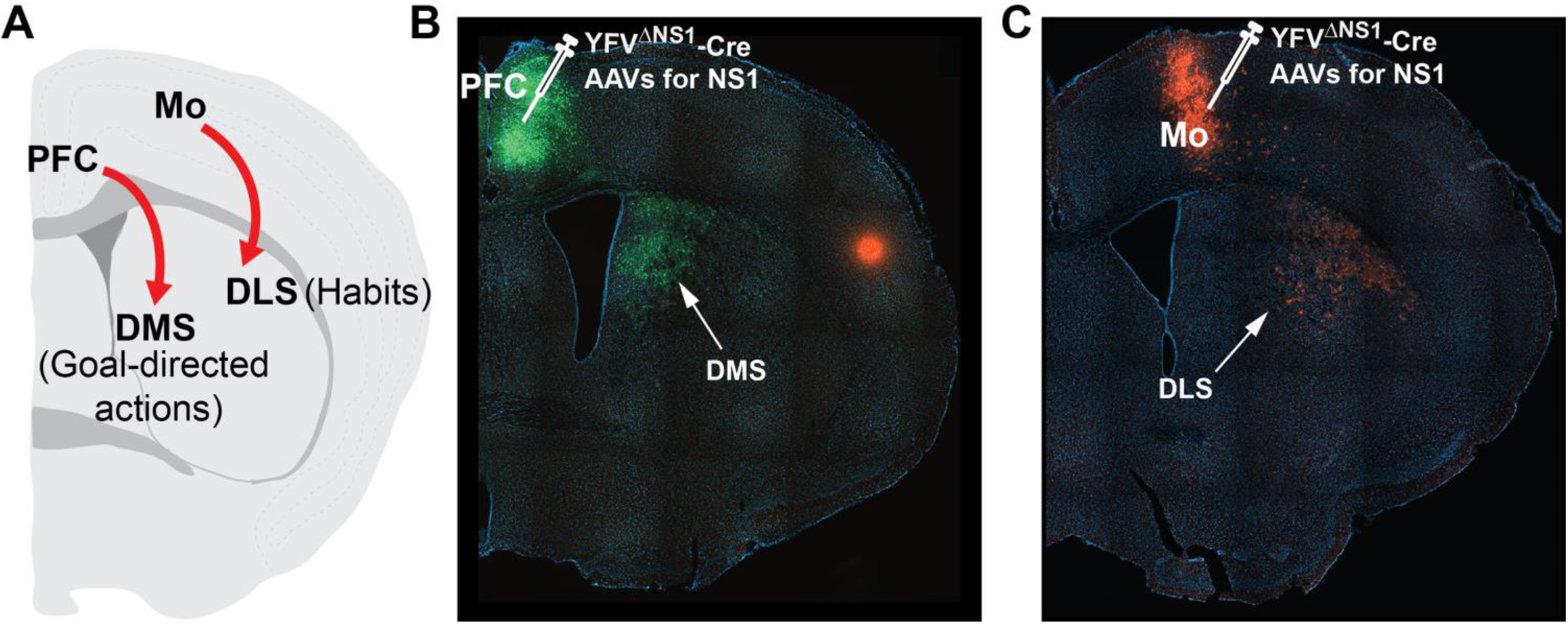
YFV^ΔNS1^-Cre traces the astrocytes associated with distinct corticostriatal pathways. (**A**) Schematics showing two parallel corticostriatal pathways involved in different behavioral modes of animals. (**B**) Expression of ZsGreen1 in the dorsomedial striatum (DMS) after YFV^ΔNS1^-Cre was injected into the PFC in an Ai6 mouse. (**C**) Expression of tdTomato in the dorsolateral striatum (DLS) after YFV^ΔNS1^-Cre was injected into the Mo in an Ai9 mouse.

We further examined the astrocytes associated with cortical projections to the dentate gyrus (DG). The molecular layer of DG is highly laminated. It can be divided into the outer, middle and inner molecular layers (OML, MML and IML), which receive synaptic inputs from the lateral entorhinal cortex (LEC), medial entorhinal cortex (MEC) and mossy cells, respectively (Fig. 4A). Consistent with previous reports, when we injected synaptoTAG AAV that expressed EGFP fused to synaptic vesicle protein synaptobrevin2 to the LEC, we observed EGFP-labeled synaptic terminals primarily in the OML (Fig. 4B, left panel). Similarly, EGFP-labeled synaptic terminals were primarily in the MML when synaptoTAG AAV was injected into the MEC (Fig. 4B, middle panel). We also injected AAV mediating Cre-dependent expression of hM3Dq fused to mCherry to the DG in a Calbindin2 (Calb2)-IRES-Cre mouse. The mossy cells in the DG are Calb2-positive and therefore expressed mCherry. mCherry was also present in the axonal projections from the mossy cells to the IML (Fig. 4B, right panel). We then examined the astrocytes traced from LEC or MEC, respectively. When we injected YFV^ΔNS1^-Cre to the LEC, the labeled astrocytes were in the OML (Fig. 4C); and when YFV^ΔNS1^-Cre was injected to the MEC, the labeled astrocytes were mainly in the MML, although some astrocytes were in the IML with part of their processes inside or touching the MML (Fig. 4D). These results further demonstrate that the traced astrocytes are associated with the neuronal circuits that the YFV-17D vector traced.

**Fig.4.**
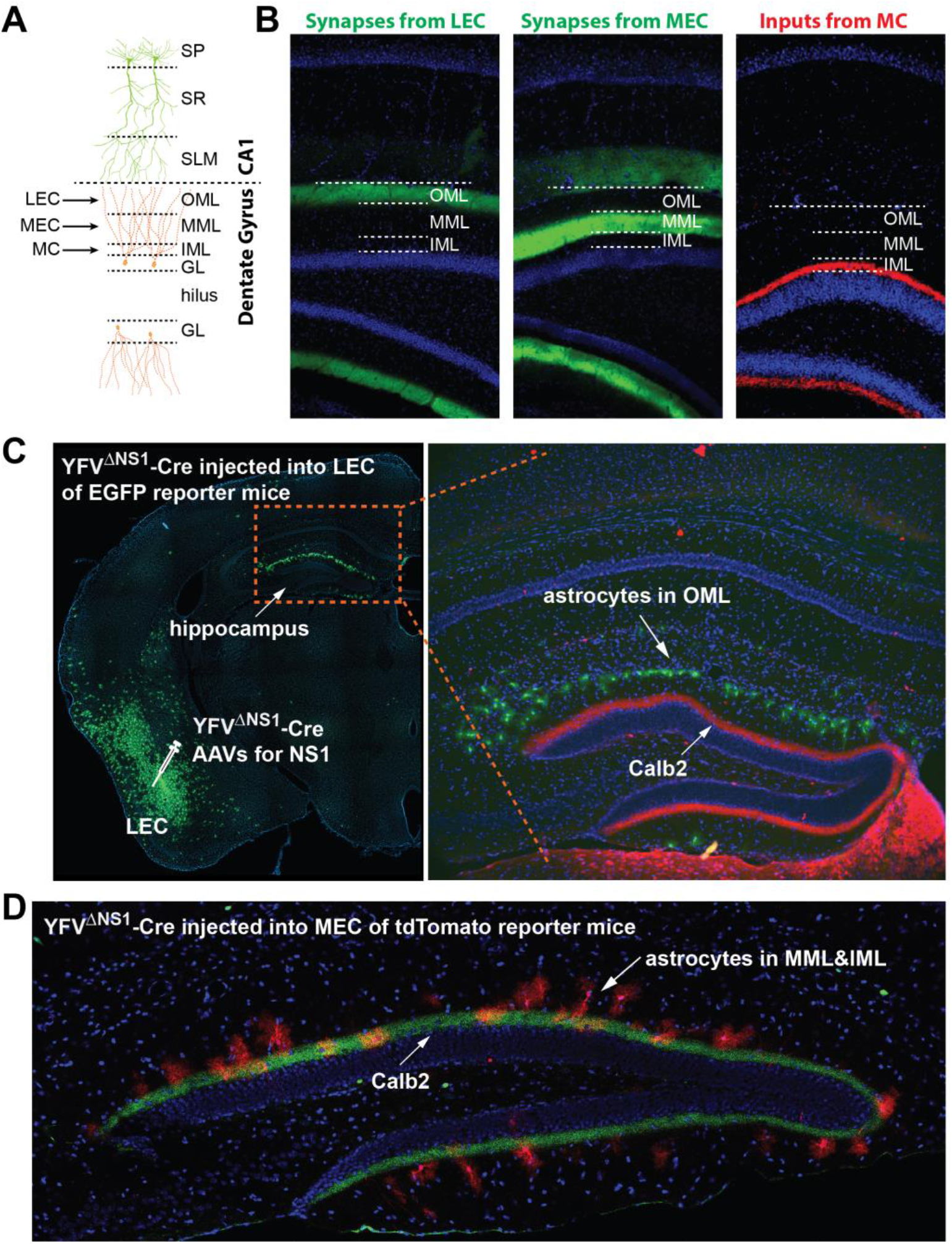
YFV^ΔNS1^-Cre traced the astrocytes associated with distinct cortex-DG pathways. (**A**) Schematics showing that three streams of axonal inputs, from LEC, MEC and MC, form synapses at the different layers of the molecular layers of the DG--the OML, MML and IML--respectively. (**B**) Expression of EGFP fused to synaptobrevin-2 after SynaptoTAG AAV was injected into the LEC (left panel) or MEC (middle panel) respectively. The right panel shows the expression of mCherry in IML of DG after injection of Cre-dependent AAV expressing hM3Dq fused with mCherry into the DG (contralateral to the hemisphere the photo was taken) in a Calb2-Cre mouse. (**C**) Expression of ZsGreen1 by astrocytes in the OML after YFV^ΔNS1^-Cre was injected into the LEC in an Ai6 mouse. The brain sections were immunostained with Calb2 (red) to visualize the axons from MC. (**D**) Expression of tdTomato by astrocytes in the MML and IML after YFV^ΔNS1^-Cre was injected into the MEC in an Ai9 mouse. The brain sections were immunostained with Calb2 (green).

## Discussion

Although astrocytes are active players of neuronal circuits, due to a lack of tools, astrocytes’ circuit-specific roles are just starting to be uncovered. Numerous novel technologies have recently been developed to target, monitor or manipulate neurons, but the tools for studying glial cells are scanty. Since some serotypes of AAVs can infect astrocytes, they can be combined with astrocyte-specific promoters or transgenic mouse lines to study astrocytes in chosen brain regions (Chai et al., 2017). Although a few AAV serotypes can travel along the neuronal circuits in anterograde or retrograde direction (Card et al., 1993; Zingg et al., 2020), no AAV has been found to efficiently target the astrocytes associated with the circuits it traces. Here we show that YFV-17D vectors carrying Cre provide this much-needed technology.

Previously we used YFV-17D vectors carrying fluorescent proteins (mVenus or mCherry) or Cre to track neuronal projections. For the replication-deficient versions, such as the YFV^ΔNS1^-mVenus or YFV^ΔNS1^-Cre, we needed to express the NS1 gene in the post-synaptic neurons to efficiently reveal these neurons. Without post-synaptic expression of the NS1 gene, the labeling would be very sparse (in the case of YFV^ΔNS1^-Cre) or non-detectible (in the case of YFV^ΔNS1^-mVenus) (Li et al., 2021). Why did these vectors effectively label astrocytes in the reporter mice? It is possible that these YFV-17D vectors can infect astrocytes but lack the ability to replicate in astrocytes. An astrocyte is frequently associated with 100,000 synapses, an order of magnitude higher than that of most neurons (Bushong et al., 2002). In reporter mice, through the large number of synapses they are associated with, each astrocyte may get multiple copies of the YFV-17D vector, which could express Cre to a sufficient level to turn on the reporter gene before the viral RNA (YFV-17D is a RNA virus) is degraded. However, due to the lack of replication, these vectors cannot amplify themselves and express mVenus to a detectible level.

With this set of tools, a few types of experiments can be conducted to understand the circuit-specific roles of astrocytes. By using these tools in reporter mice for fluorescent proteins or calcium indicators (Daigle et al., 2018), we can examine the morphological or activity features of the astrocytes distributed in the same brain region but associated with distinct neuronal circuits. By applying them in reporter mice for DREADDs or channelrhodopsin (Zhu et al., 2016), we can selectively stimulate these astrocytes with high temporal and spatial precision. We may also apply them in various conditional knockout mouse lines to determine the contributions of proteins expressed by astrocytes to synapse formation or function (Stogsdill et al., 2017).

This set of tools does have limitations. Cre may turn on the reporters in some neurons at the post-synaptic regions, albeit they are a small fraction of the cells labeled. This can be solved by placing the reporter genes under the control of astrocyte-specific promoters, but it may involve significant work to generate the reporter mouse lines. Using AAV reporter under astrocyte-specific promoters could be a convenient alternative. However, in our preliminary tests, we were not able to efficiently turn on an AAV reporter in the post-synaptic brain regions. Potential solutions to this problem may require some protocol adjustment, like elongating the time window between reporter AAV injection and YFV-17D injection, or pharmacological treatments to reduce cell antiviral functions. “Neurotrophic” viruses previously used solely for tracing neuronal circuits, like the recombinant rabies virus, can infect non-neuronal cells in the brain as well (Clark et al., 2021). Similar findings were made with pseudorabies virus (PRV) and herpes simplex virus 1 (HSV-1) (Card et al., 1993; McCarthy et al., 1990). Engineering them may further expand the tool set for glial functional analysis.

In line with their roles in synaptic functions, astrocytes are critically involved in the pathogenesis of multiple brain disorders, such as amyotrophic lateral sclerosis (Izrael et al., 2020), Alzheimer’s disease (Gonzalez-Reyes et al., 2017), autism spectrum disorder (Petrelli et al., 2016), epilepsy (Coulter and Steinhauser, 2015), and so on. The astrocytes contributing to a disorder may be located in various regions but may also be associated with certain neuronal pathways (for instance, in the case of ALS, astrocytes related to the motor cortex-spinal cord circuit may contribute to the disorder). The lack of technology to target astrocytes in a pathway-specific manner impedes the development of effective therapies. Therefore, in addition to in-depth analysis of the circuit-specific functions of astrocytes, this new technology may also be developed into treatments for neuropsychiatric disorders in which astrocytes play a central role.

## Methods

### Animals

Male tdTomato reporter mice (Ai9 mice, JAX stock No. 007909) and ZsGreen1 reporter mice (Ai6 mice, JAX stock No. 007906), and Calb2-IRES-Cre mice (JAX stock No. 010774) were group housed on a 12 hr light /12 hr dark cycle with ad libitum access to food and water. Mice were randomly assigned to experimental groups. Animal work was approved and conducted under the oversight of the UT Southwestern Institutional Animal Care and Use Committee and complied with Guide for the Care and Use of Laboratory Animals by National Research Council.

### Preparation of viral vectors

AAVs were packaged with AAV-DJ or AAV5 capsids. Virus was prepared as described (Zolotukhin et al., 1999). Briefly, AAV vectors were co-transfected with pHelper and pRC-DJ or rAAV2-retro helper into HEK293 cells. Cells were collected 72 hr later, lysed, and loaded onto iodixanol gradient for centrifugation at 400,000 g for 2 hr. The fraction with 40% iodixanol of the gradient was collected, washed, and concentrated with 100,000 MWCO tube filter. The genomic titer of virus was measured with quantitative real-time PCR. The titers of AAVs used for stereotaxic injection were in the range of 0.5-2 x 10^13^ genome copies/ml.

To generate YFV-17D stocks, plasmids were linearized with XhoI digestion and transcribed with an SP6 in vitro transcription kit (mMESSAGE mMACHINE Kit, Ambion). In vitro transcribed viral RNAs, generated from linearized plasmids encoding YFV^ΔNS1^-Cre, was transfected into BHK cells with TransIT-mRNA Transfection Kit (Mirus Bio, Cat. No. MIR 2225). Virus containing supernatant was concentrated, aliquoted, stored at −80°C, and titered by quantitative real-time PCR. The titers of YFV^ΔNS1^-Cre was 9.60X10^10^ genome copies/ml.

### Stereotaxic injections

Mice were anesthetized with tribromoethanol (125-250 mg/kg) or isoflurane inhalation (1.5-2%). Viral solution was injected with a glass pipette at a flow rate of 0.10 μl/min. After the completion of injection, the glass pipette was left in place for 5 min before being retrieved slowly. We unilaterally injected between 0.25 – 0.5 μl of viral solution at each injection site unless stated otherwise. The coordinates for each of the injection sites on the anterior-posterior direction from the bregma (AP), medial-lateral direction from the midline, and the dorsal-ventral direction from the dura (DV) are as the following: Mo (+0.45, 1.50. 0.80), PFC (+1.25, 0.30, 1.25); LEC (−3.20, 4.60, 3.60); MEC (−4.80, 3.45, 2.75) (In the AP direction, “+” denotes anterior to the bregma and “-” denotes posterior to bregma).

### Histology and immunohistochemistry

Mice were transcardially perfused with 10 ml of PBS followed by 30-40 ml of 4% paraformaldehyde (PFA) in PBS. The brains were extracted and postfixed overnight in 4% PFA at 4°C, and cryoprotected in 30% sucrose. Brains were sectioned with a cryostat to a thickness of 30 or 40-μm. Free-floating sections were washed in PBS, incubated with DAPI (1μg/ml) and mounted on slides. For immunohistochemistry, the brain sections were washed in PBS three times and permeabilized in 2% triton X-100 in PBS for 2 h. The sections were then incubated for 2 h in PBS containing 10% horse serum, 0.2% bovine serum albumin and 0.5% triton X-100, followed by overnight incubation in primary antibodies diluted in PBS containing 1% horse serum and 0.2% bovine serum albumin. The primary antibodies used and their dilutions were: NeuN antibody (clone A60), Sigma MAB377, 1:1,000; S100β, Abcam ab41548, 1:500; TMEM119 antibody, AbCam ab209064, 1:500; NG2 antibody, Sigma/Millipore AB5320, 1:500. After three washes in PBS, the sections were then incubated in the secondary antibodies for 2 h, washed again and then mounted onto glass slides. The secondary antibodies used and their dilutions were Alexa Fluor 594-conjugated goat anti-mouse (Thermo Fisher A-11032) or goat anti-rabbit IgG (Thermo Fisher A-11012) 1:500. The whole-mount brain sections were scanned with Zeiss AxioscanZ1 digital slide scanner with a 10X objective. The high-resolution images were taken with ZEISS LSM 880 with Airyscan confocal microscope.

## Acknowledgements

We thank Elizabeth Li, Heankel Oliveros, Jun Guo and So Jung Oh for contribution at the early stage of this project. This project was supported by a grant from NIH/NIMH (MH099153 to WX). We also thank Dr. Denise Ramirez (Whole Brain Microscopy Facility at UT Southwestern) and Dr. Shin Yamazaki (Neuroscience Microscopy Facility at UT Southwestern) for help with imaging.

